# The piRNA Response to Retroviral Invasion of the Koala Genome

**DOI:** 10.1101/599852

**Authors:** Tianxiong Yu, Birgit S. Koppetsch, Keith Chappell, Sara Pagliarani, Stephen Johnston, Zhiping Weng, William E. Theurkauf

**Affiliations:** Department of Clinical Laboratory Medicine, Shanghai Tenth People’s Hospital of Tongji University, The School of Life Sciences and Technology, Tongji University, Shanghai 200092, China; Program in Bioinformatics and Integrative Biology, University of Massachusetts Medical School, Worcester, MA 01605, USA; Program in Molecular Medicine, University of Massachusetts Medical School, Worcester, MA 01605, USA; School of Chemistry and Molecular Biosciences, University of Queensland, Brisbane, Australia; School of Agriculture and Food Sciences, University of Queensland, Gatton, Australia

## Abstract

Transposons are ubiquitous mobile elements with the potential to trigger genome instability and mutations linked to diseases^1,2^. Antisense piRNAs guide an adaptive genome immune system that silences established transposons during germline development^3^, but how the germline responds to new genome invaders is not understood. The KoRV retrovirus infects somatic and germline cells and is sweeping through wild koala populations by a combination of horizontal and vertical transfers, providing a unique opportunity to directly analyze the germline response to retroviral invasions of a mammalian genome^4,5^. We analyzed genome organization and long RNA and short RNA transcriptomes in testis, liver, and brain from two wild koalas infected with KoRV, while integrating our results with earlier genomic data. Consistent with data from other mammals^6,7^, koala piRNAs were detected in testis and mapped to both isolated transposon insertions and genic and intergenic piRNA clusters. Established transposon subfamilies produced roughly equal levels of antisense piRNAs, which are the effectors of trans-silencing, and sense piRNAs, which drive ping-pong amplification of these effectors^8,9^. KoRV piRNAs, in striking contrast, were strongly sense biased in both animals analyzed. These two koalas each carried 60 germline KoRV-A insertions, but only 14 of the insertions were shared, and none of the insertions mapped to piRNA clusters. The sense piRNAs thus appear to be produced by direct processing of the transcripts from isolated proviral insertions. A typical gammaretrovirus, KoRV produces spliced *Env* mRNAs and unspliced transcripts encoding Gag, Pol, and the viral genome. KoRV *Env* mRNAs were 5-fold more abundant than the unspliced pre-mRNAs, but 92% of piRNAs were derived from the unspliced pre-mRNAs. We show that this biased piRNA production from unspliced retrotransposon transcripts is conserved from flies to mice. Retroviruses must bypass splicing to replicate; thus, we propose that failed splicing produces a “molecular pattern” on transcripts from retroviral invaders that is recognized by an innate genome immune system, which silences transposons in cis by processing their transcripts into piRNAs. This innate immune response defends the germline until antisense piRNA production—from clusters or isolated insertions—is established to provide sequence-specific adaptive immunity and memory of the genome invader.

---

The 23-35 nucleotide (nt) long piRNAs bind to the PIWI-clade of Argonaute proteins and guide trans-silencing through base pairing to target RNAs, which leads to transcript destruction and post-transcriptional silencing, or inhibitory chromatin modifications and transcriptional silencing^6^. Transposon silencing by the piRNA pathway is best understood in *Drosophila*, where the majority of these small RNAs are derived from long non-coding genomic “clusters”, composed of nested transposon fragments^9–11^. The *Drosophila* piRNA pathway has distinct germline and somatic branches. Germline clusters produce piRNAs from both genomic strands, albeit with an antisense bias (with respect to transposon mRNAs), and contain randomly oriented transposon fragments. By contrast, in the dominant somatic cluster, *flamenco* (*flam*), transposon insertions are primarily in one orientation, piRNAs are strongly antisense biased, and *flam* mutations lead to sterility and over-expression of those transposons represented in the locus^12,13^. The majority of piRNA clusters in mice and other mammals are also transcribed from one genomic strand^14^, and transposon insertions in mouse clusters are antisense biased^15^. These findings suggest that transposon insertion into clusters generates antisense piRNAs, which guide sequence-specific adaptive genome immunity.

Analysis of hybrid dysgenesis in *Drosophila* has provided insights into piRNA pathway adaptation to DNA transposon invasion in insects^16–18^, but the response to invasions of the mammalian genome has not been examined. KoRV-A is associated with leukemia, immunodeficiency and chlamydia infection in koala, and characterization of proviral integration led to the surprising discovery that the virus had entered the koala germline in the last 50,000 years and is spreading by an unusual combination of vertical and horizontal transfers^4,5,19,20^. The virus is believed to have entered the wild population at the northern end of Australia and swept south, where a few naive populations persist^5^. Koala infection by KoRV-A thus provides a unique opportunity to directly examine the progression of the germline response to retroviral invasions in a wild population.

## KoRV-A and three endogenous retrotransposons are active in the koala germline

To define the genomic context of KoRV-A invasion, we characterized endogenous transposons in the Koala reference genome (see Methods). This analysis identified 402 transposon subfamilies, which occupy 44.04% of the genome. Most of the transposon subfamilies are represented by short, degenerate copies, which appear to be inactive (Fig. S1a, gray points). However, the reference genome carries seven full-length copies of KoRV-A with less than 0.5% sequence divergence (Fig. S1a), confirming that it has been active recently^5,21,22^. The reference genome also carries multiple, full-length copies of three endogenous retroviruses, designated ERVL.1, ERV.1, and ERVK.14, which show less than 1% divergence (Fig. S1a, as indicated), suggesting that these endogenous elements may also be active.

To assess expression of the transposon subfamilies, we examined the transcriptomes in a panel of tissues from four wild koalas, by performing RNA-sequencing (RNA-seq) on two previously uncharacterized individuals (designated K63464 and K63855) and reanalyzing the published RNA-seq data of two additional individuals (Birke and PC^23^). Only the handful of transposons with full-length copies were highly expressed in multiple tissues, including testis, with KoRV-A showing the highest expression levels, followed by ERVL.1, ERV.1, and ERVK.14 (Fig. 1a). Elevated transcription of KoRV-A and the three endogenous retrotransposons suggests that they all are currently active.

**Figure 1.**
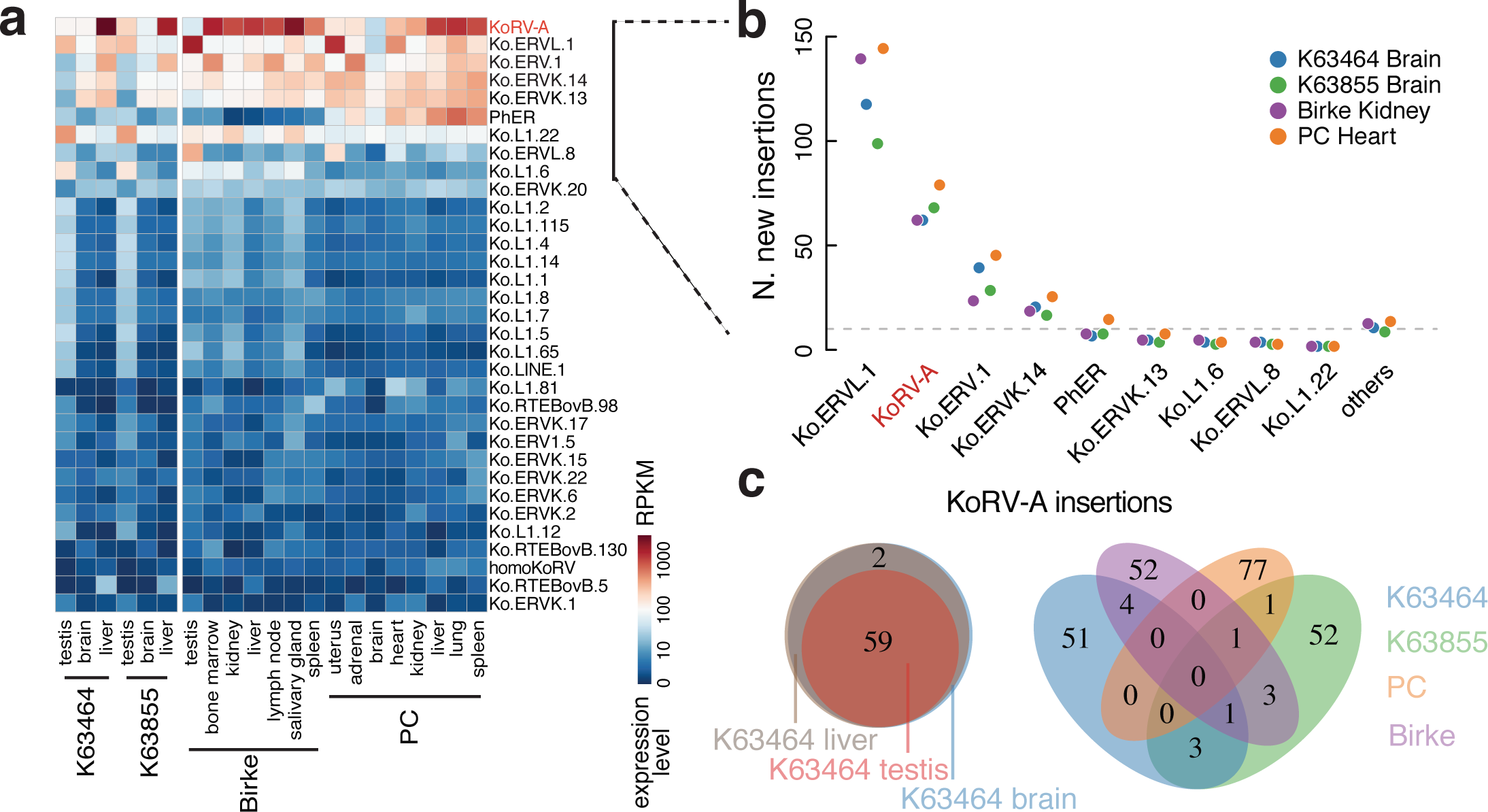
KoRV-A is highly expressed and actively transposing in the koala genome. **a**. heatmap depicts the expression levels of transposon subfamilies in tissues from four koalas: K63464 and K63855 this work and Birke and PC^21,23^. Only the transposons with expression levels higher than 1 RPKM are shown. **b**. Number of new germline insertions (defined as having reads mapped to both ends of the integration site and penetrance greater than 0.4 in at least one tissue from a koala individual). The nine most highly expressed transposon subfamilies from **Figure 1a** are graphed individually, and the remaining transposon subfamilies are pooled and labeled as others. **c.** Venn diagrams show the overlap of KoRV-A insertions among tissue samples. (Left) The number of shared KoRV-A insertions in the testis, liver and brain genomes of K63464. (Right) Shared KoRV-A germline insertions in K63464, K63855, Birke, and PC genomes.

Ongoing germline transposition would generate individual-to-individual variation, while recent activity in a common ancestor would produce low-divergence insertions that are shared between individuals. Previous analyses indicate that KoRV is currently active and generating significant individual-to-individual variation^5,24^. To extend these earlier studies and assay for endogenous retrotransposon activity, we directly identified transposon insertion sites in the four wild koalas, by sequencing genomic DNA from K63464 and K63855 and analyzing published genome sequences from Birke and PC^21^. We mapped new transposon insertions—insertions not in the reference koala genome (the genome of a koala named Bilbo^21^)—and quantified their insertion frequencies as described in Methods. To avoid low-frequency chimeric reads that result from sequencing library construction^25^, we required both ends of the integration site of each insertion to be supported by “discordant” read pairs, with one read mapping to the reference genome and the other read of the pair mapping to the inserted element (see Methods). Indeed, consistent with earlier studies, we identified new insertions for KoRV-A, but also identified numerous new insertions of the three endogenous retrotransposons (Fig. 1b), confirming that all of these elements are active and altering the genomes of these koalas.

We next defined germline integration by new insertions present at 40% or greater frequency (consistent with heterozygosity) in at least one tissue. For K63464, we sequenced genomic DNA from testis, liver, and brain, and for K63855, we sequenced genomic DNA from liver and brain. For these two animals, we further required germline insertions to be present in all of the tissues analyzed (Fig. 1c, Fig. S1b). By these criteria, the four animals carried ∼60 germline KoRV-A insertions; however, none of the germline insertions was shared by all four individuals: Birke and PC shared a single KoRV-A insertion, while K63464 and K63855 shared foursites (Fig. 1c). These four animals, which were all collected from the center of the habitat range, thus are likely to represent at least two independent KoRV-A genome invasion events, followed by significant KoRV-A expansion in the germline. In contrast, we identified 22 ERVL.1, 15 ERV.1, and 2 ERVK.14 germline insertions that were shared by all four animals (Fig. S1c), confirming that these established endogenous retrotransposon subfamilies were active in a common ancestor. However, each animal also carried numerous unique insertions of each of these elements (Fig. S1c). For example, K63855 had eight germline ERVK.14 insertions that were not present in any other animals, and PC had 94 unique germline ERVL.1 insertions (Fig. S1c). KoRV-A and the three endogenous retrotransposons are therefore generating remarkable germline variations in wild koalas.

To determine if KoRV-A and endogenous retrotransposons are active during development or in adult tissues, we quantified *de novo* transposon insertions, defined as insertions only present in one tissue and supported by only one pair of discordant reads (see Methods). Some discordant read pairs arise from low-frequency sequencing artifacts that randomly join genomic DNA fragments^25^, but the frequency of such read pairs should be constant across different tissues from the same animal, which share a common genome, and the transposon-mapping ends of these artificial reads should be randomly distributed over the element. Therefore, we analyzed genomic DNA sequences from the testis, brain, and liver of a single animal, K63464, and filtered for reads mapping to terminal regions of mobile elements. This analysis revealed different apparent transposition rates in the three tissues, with the lowest rate in testis, followed by brain and liver (Fig. S1d). KoRV-A also showed strikingly elevated numbers of discordant reads in the liver, and RNA-seq data indicate that KoRV-A is overexpressed in this tissue (Fig. 1a). These findings are likely to be related to the proportionately higher level of blood leukocytes within this tissue. While our sample size is extremely limited, these results suggest that KoRV-A and the endogenous retrotransposons are active in both the germline and the soma, and their activities are relatively low in the testis, where the piRNA pathway operates.

## Genomic sources of koala piRNAs

To determine if piRNAs are produced to target the active KoRV-A and endogenous retrotransposons, we sequenced 18–34 nt long small RNAs from testis, liver, and brain, for both K63464 and K63855 (piRNAs are typically 24–32 nt long). Furthermore, because piRNAs are 2’– O-methylated at their 3’ termini, which renders them resistant to oxidation, we sequenced both unoxidized and oxidized libraries for each koala. We detected 24–32 nt, oxidation-resistant RNAs only in the testis (Fig. S3a, b), and these RNAs showed a strong 5’-U (1U) bias and significant 10-nt overlaps between sense and antisense species (ping-pong signature, Fig. S2)—hallmarks of piRNAs observed in previously characterized animals, including flies and mice^9,26–28^. Accordingly, most of the 19 genes annotated to function in the piRNA pathway are more highly expressed in the testis than in somatic tissues (Fig. S3).

In the mammals analyzed to date, piRNAs are produced from long non-coding intergenic clusters, genic clusters that encode proteins expressed in the soma and piRNAs processed in the germline, and isolated transposon insertions^14,29,30^. Intergenic and genic clusters can be single transcription units or divergently transcribed pairs of transcription units. Mapping piRNAs and long RNA reads in the K63464 testis to the reference koala genome (see Methods) revealed the same spectrum of piRNA clusters (Fig. 2a-f). In total, we annotated 376 piRNA clusters, with the vast majority of them producing piRNAs from only one genomic strand, including 188 unidirectional clusters in protein-coding genic loci, 126 unidirectional clusters in intergenic loci, and 36 bidirectional clusters in intergenic loci (Fig. 2g). Thus, piRNAs and long RNAs in clusters map predominantly to the same genomic strand (Fig. 2h, S5a), and the piRNAs exhibit a strong 1U bias and ping-pong signature (Fig. 2i-j). The promoters of the intergenic unidirectional and bidirectional piRNA clusters are enriched for the binding motif of A-MYB (Fig. S4a), a master regulator of pachytene piRNA production in mice^14^. RNA sequencing indicates that the 162 intergenic piRNA clusters are transcribed specifically in the testis (Fig. S4b), while the genic loci are transcribed in all three tissues (Fig. S4b). Cumulatively, these 376 clusters occupy only 0.17% of the koala genome but can account for 68.53% of all piRNAs (75.09% of unique-mapping piRNAs and up to 46.56% of multiple-mapping piRNAs). Isolated transposon insertions, which occupy 43.99% of the genome, can account for up to 32.09% of all piRNAs (18.2% of piRNAs map to both clusters and dispersed transposon copies). Transposon-mapping piRNAs are abundant during pre-pachytene stages when the piRNA pathway is required for transposon silencing; however, unique piRNAs from highly expressed pachytene clusters dominate the piRNA pool in adult mouse testes^14^. Adult koala testes likely show a similar temporal expression pattern of pachytene clusters. Koalas are therefore similar to other mammals and produce piRNAs from a combination of clusters and isolated transposon insertions.

**Figure 2.**
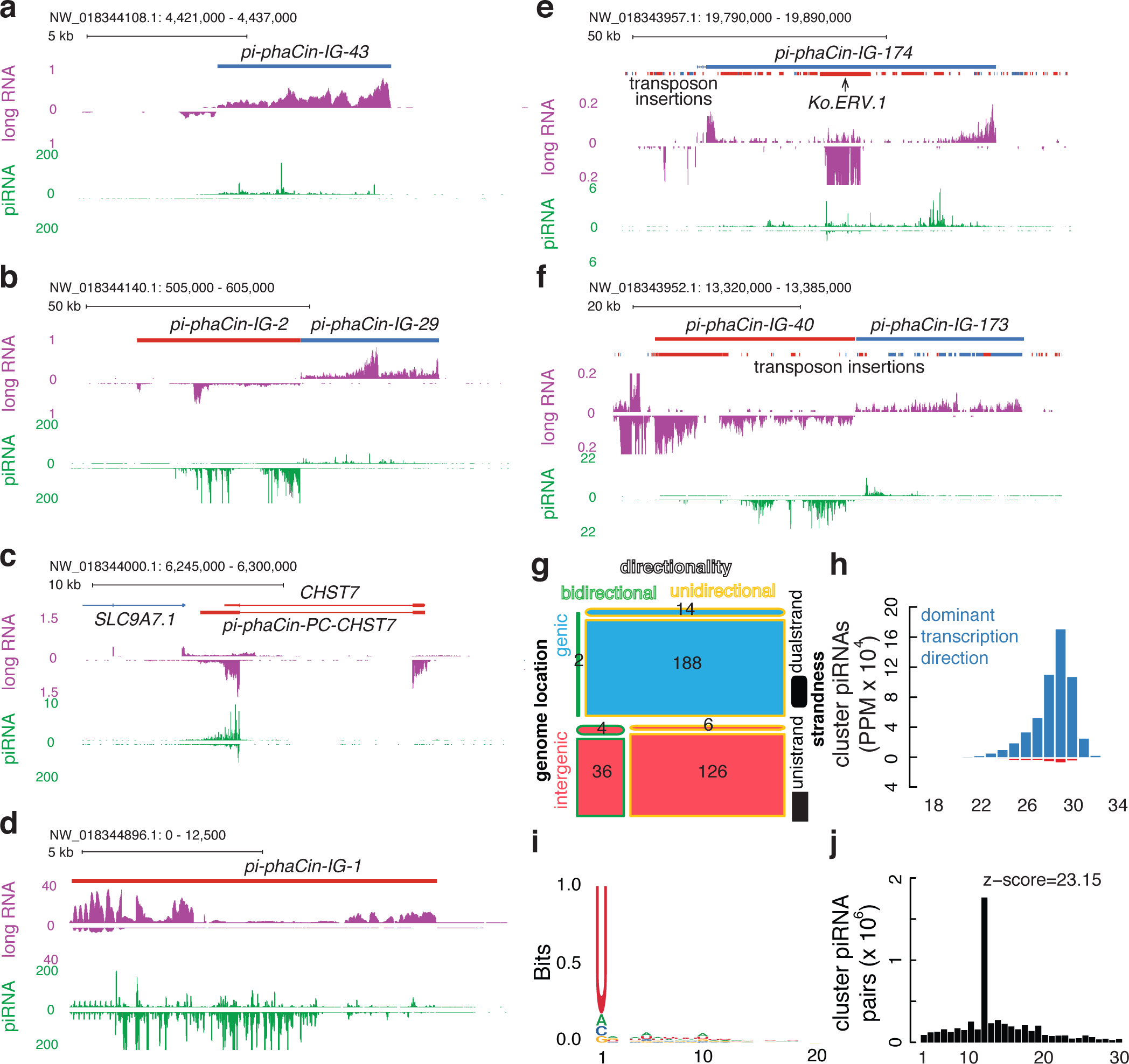
Annotation of piRNA clusters in the koala genome. **a.** An intergenic, unidirectional, and uni-strand piRNA cluster. The top tracks show the location of the piRNA cluster, gene annotation, and transposon insertions, where appropriate. Plus-strand piRNA clusters, genes, or transposon copies are shown in blue while minus-strand copies are in red. The middle track in purple shows long-RNA abundance (in RPKM) and the bottom track in green shows piRNA abundance (in PPM). These track notations are also used in **b**-**f**. **b.** A pair of intergenic, bidirectional, and uni-strand piRNA clusters. **c.** A genic, unidirectional, and dual-strand piRNA cluster producing piRNAs from the overlapping 3’-UTRs of two convergently transcribed protein-coding genes *SLC9A7.1* and *CHST7*. **d.** An intergenic, unidirectional, and dual-strand piRNA cluster. **e.** An intergenic, unidirectional, and uni-strand piRNA cluster in the plus strand with a Ko.ERV.1 inserted in the minus strand. **f.** A pair of intergenic, bidirectional, and uni-strand piRNA clusters with sense-biased transposon insertions. **g.** A mosaic plot for 376 piRNA clusters classified by: (1) genomic location (genic in blue, characterized by the overlap of piRNA clusters with protein-coding genes, and intergenic in red); (2) directionality (bidirectional, divergently transcribed piRNA clusters are shown with a green outline and unidirectional piRNA clusters in yellow); and (3) strandedness (dual-strand if the ratio of piRNAs abundance from the two genomic strands are within four folds, depicted as obrounds, and uni-strand as rectangles). Black numbers denote the tallies of piRNA clusters in each classification; there are zero bidirectional, genic piRNA clusters. **h.** Size distribution of piRNAs derived from piRNA clusters in the oxidized K63855 testis. **i**. A sequence logo shows the nucleotide composition of the piRNAs derived from piRNA clusters in the oxidized K63855 testis. **j.** Distribution of overlapping nucleotides between plus-strand and minus-strand piRNAs derived from piRNA clusters in the oxidized K63855 testis. The prominent peak at the 10-nt overlap is the ping-pong signature.

## Are transposons with cluster insertions more likely to produce antisense piRNAs?

Roughly equal numbers of sense and antisense piRNAs map to transposons and these piRNAs show the 1U-10A nucleotide bias and ping-pong signature (Fig. 3a-c, Fig. S5b, d). Transposon insertion into a cluster was proposed to enhance the production of trans-silencing antisense piRNAs^31^; thus, we asked whether transposon subfamilies with cluster insertions were more likely to produce antisense-biased piRNAs than subfamilies with only dispersed copies. However, regardless of piRNA cluster representation, transposon subfamilies produce comparable numbers of sense and antisense piRNAs, although there is a large variation among subfamilies (Fig. 3d-e, Fig. S5c). Transposon subfamilies with abundant piRNAs are generally represented in clusters (Fig. 3d), but these subfamilies are also more abundant in the rest of the genome than the subfamilies that produce few piRNAs (data not shown), and we did not observe a systematic difference between the two types of subfamilies in terms of bias toward antisense piRNAs. We performed the same analysis on adult mouse testis and reached the same conclusion (data not shown). Like mice, koalas piRNA clusters are slightly, but significantly, depleted of transposons compared with the rest of the genome (Fig. S4c, Wilcoxon signed-rank test p-value = 3.1 × 10^-7^ and < 2.2 × 10^-16^ for intergenic and genic piRNA clusters, respectively). Thus, our data do not support the hypothesis that transposon subfamilies with cluster insertions are in general more likely to produce antisense-biased piRNAs.

**Figure 3.**
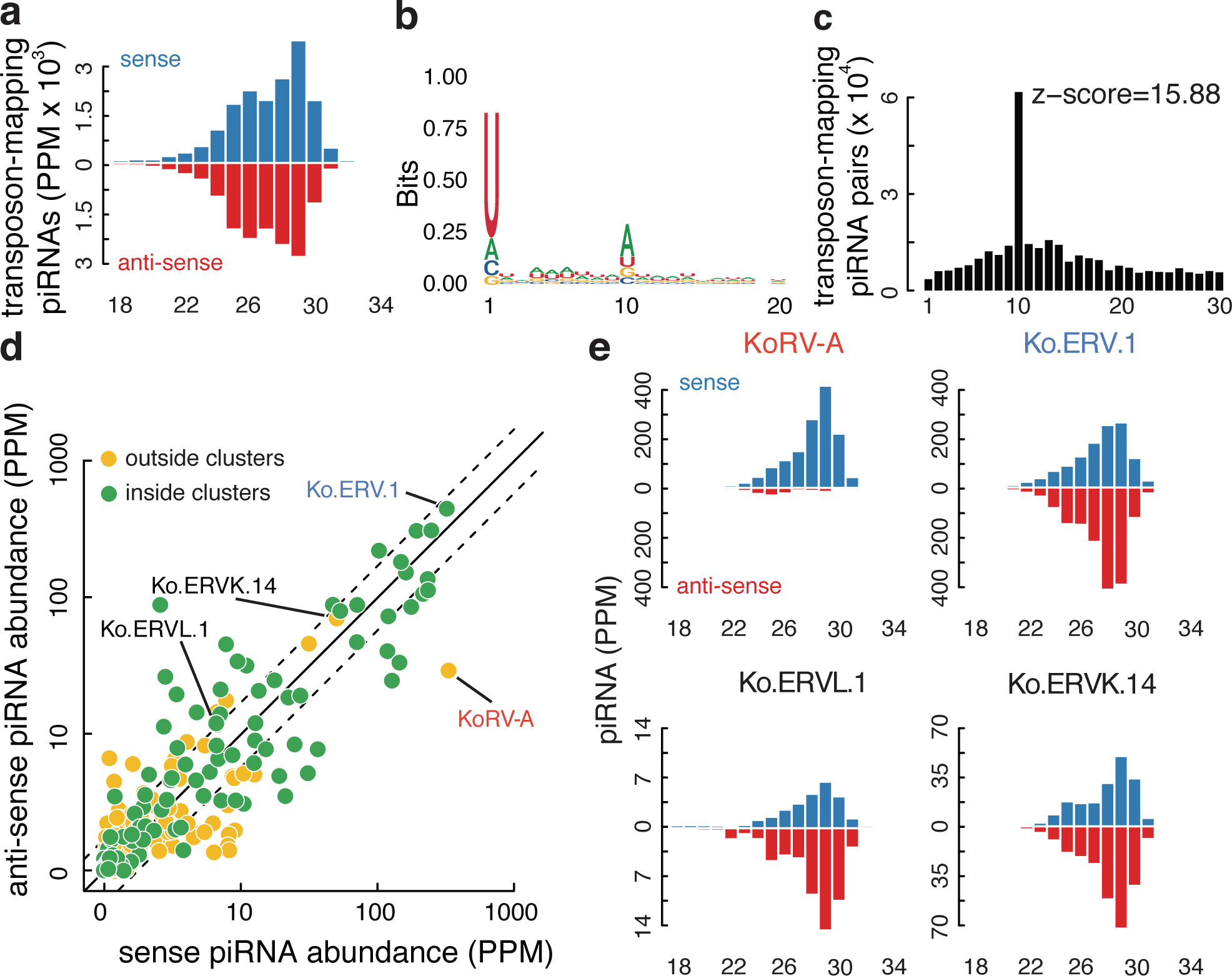
KoRV-A piRNAs are sense biased. **a.** Size distribution for small RNAs targeting all transposons in the oxidized K63855 testis. **b.** A sequence logo for transposon-targeting piRNAs in the oxidized K63855 testis. **c.** Distribution of overlapping nucleotides between sense and antisense transposon-targeting piRNAs in the oxidized K63855 testis. **d.** A scatterplot comparing the abundance (in PPM) of sense piRNAs and antisense piRNAs targeting all transposon subfamilies in the oxidized K63855 testis. Each dot represents a transposon subfamily: subfamilies not present in piRNA clusters are in yellow, while the subfamilies present in piRNA clusters are in green. Half and 2-fold are marked with dashed lines. **e.** Size distributions for piRNAs targeting KoRV-A, Ko.ERV.1, Ko.ERVL.1 and Ko.ERVK.14 in the oxidized K63855 testis.

Nevertheless, a small number of koala clusters are enriched for antisense transposon insertions and produce piRNAs with a strong antisense bias (e.g., Fig. 2e). These features were first observed for the *Drosophila flam* cluster, which has an established function in transposon silencing in the somatic follicles cells of the ovary. The three active endogenous retrotransposons in koala all produce antisense-biased piRNAs (Fig. 3e, Fig. S5c). In particular, ERV.1 produces more piRNAs than any other transposon subfamily, and there is a nearly full-length antisense ERV.1 insertion in the pi-phaCin-IG-174 intergenic cluster, which is enriched for fragments of other transposons inserted in the same orientation as ERV.1 and produces predominantly antisense piRNAs (Fig. 2e). ERV.1 is also expressed at very low levels and shows little random transposition in testes (Fig. 1a, S1d), suggesting that the antisense piRNAs are effective at repressing ERV.1.

By contrast, the piRNAs mapping to KoRV-A are strongly sense-biased (Fig. 3e, S5c, and compare histograms in Fig. 3e). KoRV-A is present in a piRNA cluster in the reference genome, but this insertion is not present in the two animals we analyzed (Fig. S4d). These animals carry over 60 germline KoRV-A insertions each, but none of these insertions map to a piRNA cluster. In *Drosophila*, a subset of single euchromatic transposon insertions are bound by the cluster-specific HP1 homolog Rhino and produce piRNAs from both genomic strands, and these “mini clusters” can be identified by the presence of piRNAs mapping to sequences flanking the insertion sites^32^. We do not detect piRNAs in sequences flanking the KoRV-A insertions in two koalas we analyzed (Fig. S4e). With the caveat there are gaps in the koala genome assembly that might include additional piRNA clusters, these observations strongly suggest that the transcripts from dispersed KoRV-A proviral insertions, which do not function as clusters, are directly processed into sense piRNAs. Intriguingly, KoRV-A is expressed at very low levels in testis from Birke, and this animal shares the cluster insertion that is present in the reference genome (Fig. 1a, Birke). These observations suggest that the cluster insertion produced trans-silencing antisense piRNAs. Unfortunately, the testis tissue is no longer available from Birke, and we were unable to test this hypothesis.

## Unspliced KoRV-A transcripts produce sense piRNAs

KoRV-A is a gammaretrovirus; transcription of proviral insertions produces spliced transcripts that encode the Env protein and unspliced transcripts that encode Gag, Pol and the retroviral genome. In koala testes, spliced KoRV-A *Env* mRNAs are 5-fold more abundant than unspliced genomic transcripts (Fig. 4a). In sharp contrast, KoRV-A piRNAs are uniformly distributed over the viral genome, suggesting that they are derived almost exclusively from unspliced genomic transcripts (Fig. 4a). To further test this possibility, we quantified long and short RNA reads mapping to the exon-exon junction, which are specific to mature *Env* mRNAs, and to splice sites, which are only present in unspliced genomic transcripts. For long RNAs, the exon-exon junction to splice site ratio was 5.0, consistent with the abundance of the *Env* mRNA in testis. The ratios are lower in liver and brain (Fig. S6a), suggesting the spliced KoRV-A transcripts are mainly derived from germ cells but not somatic cells in testis. In sharp contrast, the junction to splice site ratio for piRNAs was 0.08 in testis (Fig. 4a). Assuming that piRNAs derived from spliced and unspliced transcripts have similar stability, the 62-fold difference in these ratios reflects the difference in piRNA processing efficiency.

**Figure 4.**
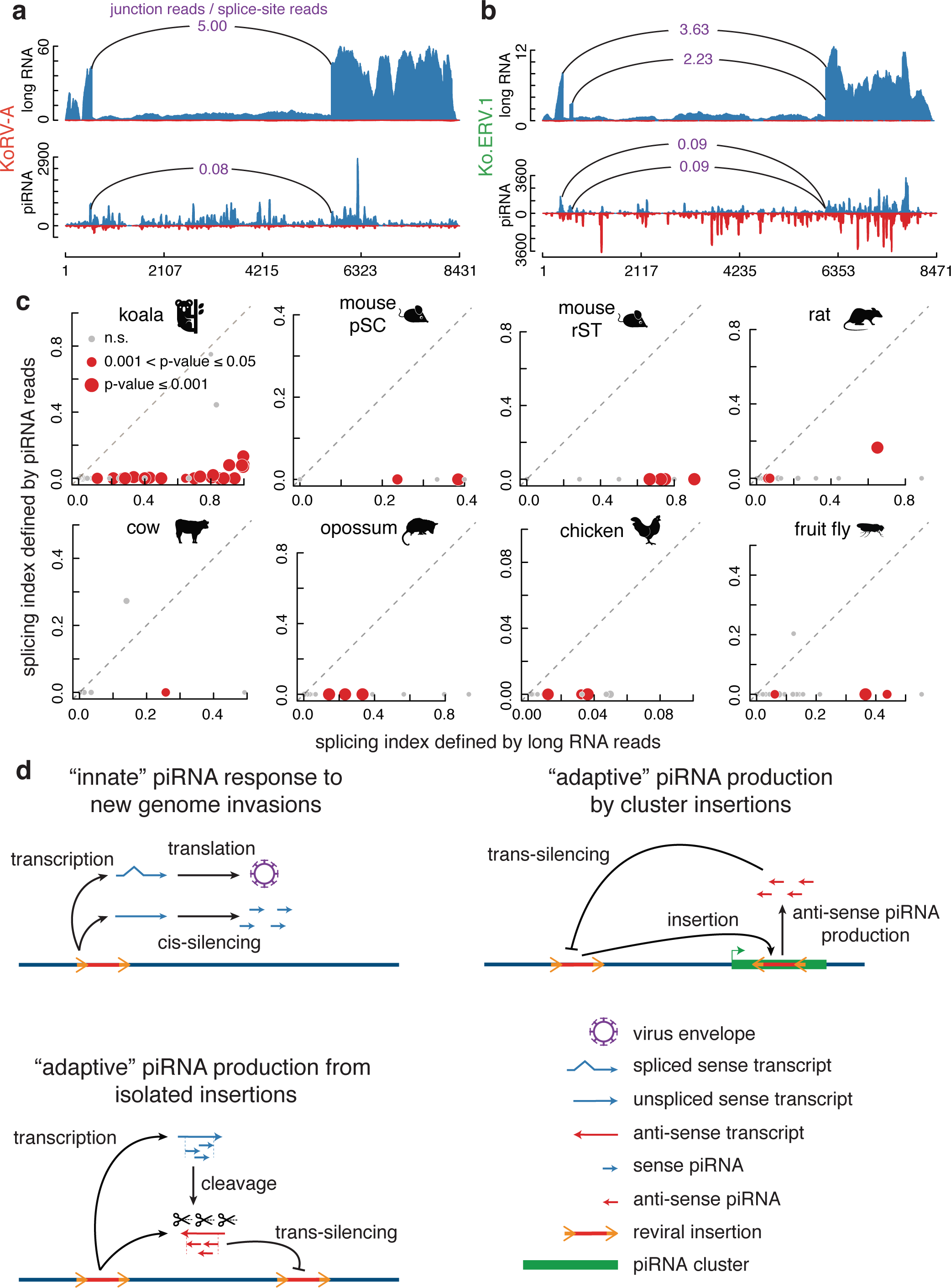
Transposon piRNAs are produced primarily from unspliced transposon transcripts. **a.** Long RNA and piRNA reads across KoRV-A in koala testis, with sense reads in blue and antisense reads in red. The junction to splice-site ratios for the main exon-exon junction is indicated, defined by long RNA or piRNA reads. **b.** Long RNA and piRNA reads across Ko.ERV.1 in koala testis, corresponding to **a** but for Ko.ERV.1 in koala. **c.** Scatterplots comparing splicing index defined by long RNA reads and piRNA reads, for all transposon exon-exon junctions in adult testes from koala, rat, cow, opossum, chicken and fruit fly, and pachytene spermatocyte (pSC) and round spermatid (rST) from mice. Exon-exon junctions with significantly higher splicing index defined by long RNA reads than by piRNA reads are colored red, otherwise in grey, and the size of the dots depict statistical significance (chi-square test p-values). **d**. Models for piRNA response to genome invasions: (top left) “innate” piRNA response to new genome invasions; (top right) “adaptive” piRNA production by insertions in piRNA clusters; (bottom left) “adaptive” piRNA production from isolated insertions.

Like KoRV-A, the endogenous ERV.1 retrotransposon showed significantly lower junction to splice-site ratios for piRNAs (0.09 for both junctions) than for long-RNAs (3.63 and 2.23), reflecting preferential processing of the unspliced genomic transcripts into piRNAs (Fig. 4b). Therefore, we asked whether preferential piRNA processing from unspliced transcripts was a general feature of koala transposons. To answer this question, we defined a “splicing index” as the ratio of exon-exon junction reads to the sum of exon-exon junction and splice site reads, with 0% indicating no reads map to the junction, while 100% indicating all reads map to the junction. For piRNAs, the splicing index was nearly 0% for all koala transposon subfamilies, but long RNA splicing index ranged from 10% to over 90% (Fig. 4c). In striking contrast, the long-RNAs from genic piRNA clusters are efficiently spliced, and the piRNAs from these clusters are preferentially produced from the spliced transcripts (Fig. S6b-c). Retrotransposon-derived and genic piRNAs thus appear to be produced by distinct mechanisms, preferentially utilizing unspliced and spliced transcripts, respectively.

We extended this analysis to other species, using published long and short RNA data from mouse, rat, cow, opossum, chicken, and fly. In each species, piRNAs were preferentially produced from unspliced sense-strand transposon transcripts (Figure 4c). For several species, including mouse, we did not detect a single piRNA read mapping to an exon-exon junction (Fig. 4c, S6d). An example is shown in Fig. S6e—the ERV1_MD_I transposon in opossum testis—with evidence of splicing in long RNAs, but piRNAs predominantly originate from unspliced transcripts. In *Drosophila*, transposon-silencing piRNAs are derived from unspliced transcripts of heterochromatic clusters composed of nested transposon fragments, and these genomic domains are bound by the Rhino-Deadlock-Cutoff complex, which is required for piRNA production by actively suppressing splicing. In addition, transposon-silencing siRNAs in pathogenic yeast are derived from stalled splicing intermediates^33^. Two components of the ubiquitous RNA-processing complex THO, Thoc5 and Thoc7, have been shown to be essential for piRNA biogenesis in flies^34^. Thoc5 and Thoc7 are required for efficient nuclear export of unspliced transcripts of murine leukemia virus in human cell lines^35^. Stalled or blocked splicing thus appears to be a conserved trigger for small silencing RNA production from potentially pathogenic elements.

### Summary

We show that KoRV-A and three endogenous retroviruses are currently active and generating significant genetic variation in wild koalas (Fig. 1, S1). Ongoing activity of the endogenous elements is surprising, as all three are targeted by antisense piRNAs (Fig. 3e). In *Drosophila*, invasion of P elements activates endogenous transposons^17^, raising the possibility that activation of endogenous retrotransposons in koala is secondary to KoRV-A invasion. The mechanism of endogenous element activation on P element invasion is not well understood, but mobilization of this DNA transposon leads to double-stranded DNA breaks, and genotoxic stress disrupts transposon silencing in a wide range of systems^36–38^. Genome instability caused by ongoing transposition of KoRV-A, which is not targeted by abundant anti-sense piRNAs, could disrupt silencing of established transposon families. Alternatively, KoRV-A, like other gammaretroviruses, encodes an immunosuppressive domain within the envelope protein (p145E)^39^, which may disrupt transposon silencing downstream of piRNA production. Both models predict that endogenous elements will be inactive in animals that have not been infected with KoRV, which are present in isolated southern populations.

Immune responses to viral or bacterial infections comprise distinct innate and adaptive phases^32^. The initial response involves recognition of common molecular “patterns” by cells of the innate immune system, which suppress invaders until a more effective adaptive response is mounted^40^. Adaptive immunity is produced by differentiation and amplification of lymphocytes, which produce antigen-specific antibodies and T cells and carry a memory of the infection^41^. The data presented here, with earlier studies, lead us to propose that the piRNA response to retroviral invasions of the genome is also composed of distinct innate and adaptive phases (Fig. 4d). During initial infection, random proviral insertions are transcribed and produce spliced and unspliced transcripts. We propose that the bypass of splicing signals generates a molecular “pattern” that differentiates genomic transcripts of spliced viral and genic mRNAs. This pattern is recognized by the innate genome immune system, which suppresses expression by directing these transcripts to the piRNA biogenesis machinery. The resulting sense-stranded piRNAs are products of this “cis-silencing” process, and not silencing effectors. As expected, somatic tissues without the “innate” piRNA “cis-silencing” system have more unspliced retroviral transcripts compare to spliced ones (Fig. S6f. splicing index of KoRV-A and ERV.1 in two koalas are shown). This system is inefficient, and retrotransposition continues until an adaptive response, mediated by antisense piRNAs, is mounted. The antisense piRNAs that guide the adaptive trans-silencing are derived from clusters carrying transposon insertions, or single insertions that are transcribed and piRNAs from both strands, through an epigenetic process that is not understood. Initiation of the adaptive genome immune response thus appears to be triggered by transposition into a cluster, or conversion of single transposon insertions into mini-clusters. Both processes generate trans-silencing effectors of adaptive immunity, while transposition into a cluster and insertion conversion produce hard-wired genetic and epigenetic memories of the genome invader.

## SUPPLEMENTARY FIGURES AND TABLES

**Figure S1. KoRV-A and three other ERVs are potentially active in the koala genome.**

**a.** A scatterplot depicts the number of full-length copies of a transposon subfamily (defined as longer than 50% of the consensus sequence length) vs. divergence from the consensus sequence. Four ERVs are labeled, including KoRV-A, which have more than five full-length copies at divergence lower than 1% in the koala genome.

**b.** Venn diagrams show the overlap of transposon insertions among the three tissues of K63464. They correspond to the left panel of **Figure 1c** but for transposon subfamilies other than KoRV-A.

**c.** Venn diagrams show the overlap of transposon insertions among the various koalas. They correspond to the right panel of **Figure 1c** but for transposon subfamilies other than KoRV-A.

**d.** Numbers of un-inherited insertions and germline insertions for all transposon subfamilies in the testis, brain, and liver of K63464. Un-inherited insertions are defined as insertions supported by only one discordant read and only present in one tissue. Germline insertions are defined as in Fig. 1b. KoRV-A, Ko.ERV.1, Ko.ERVL.1, Ko.ERVK.14 and PhER are colored in red, blue, green, yellow, and purple respectively, while other transposon subfamilies are in black.

**Figure S2. Koala piRNAs are produced in the testis but not in the brain or liver.**

**a.** Length distribution of piRNAs, nucleotide composition of piRNAs, and distribution of overlapping nucleotides between sense and antisense piRNAs. All genome-mapping oxidized small RNA reads in the K63464 testis after removal of rRNAs, miRNAs, snRNAs, snoRNAs, and tRNAs were included.

**b-f.** Correspond to panel **a** but for oxidized small RNA reads in the K63855 testis, as well as unoxidized small RNA reads in the K63464 testis, the K63855 testis, the K63464 liver, the K63855 liver, the K63464 brain, and the K63855 brain samples, respectively.

**Figure S3. Most piRNA pathway genes are specifically expressed in the koala testis.**

A heatmap showing the expression levels of 19 piRNA pathway genes in various koala tissue samples. The expression level (in RPKM) is written in each cell, and highly expressed genes are shown with three intensities of red.

**Figure S4. Koala piRNA clusters have similar features to mouse piRNA clusters**

**a.** Boxplots show the maximal A-MYB motif scores in the ±100 bp window centered on the TSSs of 19,945 mRNAs or three classes of piRNA clusters: 202 genic unidirectional, 132 intergenic unidirectional, and 42 bidirectional piRNA clusters.

**b.** Long RNA expression levels at intergenic piRNA clusters (red) and genic piRNA clusters (blue) in the testis, brain and liver samples of the two koalas assayed by us.

**c.** piRNA clusters are significantly depleted of transposons. The observed number of transposon-overlapping nucleotides in each of 376 piRNA clusters is plotted against the expected number of transposon-overlapping nucleotides. Wilcoxon rank-sum test p-value = 3.1 × 10^-7^ and < 2.2 × 10^-16^ for intergenic (red) and genic (blue) piRNA clusters, respectively.

**d.** A genic, unidirectional and uni-strand piRNA cluster with a KoRV-A insertion in the reference koala genome but the insertion is absent in our animals. The top track indicates this piRNA cluster and the KoRV-A insertion annotated in the reference genome. The middle and bottom tracks illustrate DNA-seq reads mapping around the KoRV-A insertion in K63464 and K63855 brain samples. Note the absence of reads that map to the KoRV-A despite plentiful reads in flanking regions. Only the two ends of the KoRV-A insertion are shown and the internal segment is illustrated with dashed lines (not to scale).

**e.** Profiles showing average piRNA abundance around 5 kb long KoRV-A insertions in K63464 and K63855. piRNA abundance in KoRV-A is shown in the center. piRNAs in the same strand as KoRV-A insertions are in blue while piRNAs in the opposite strand in red.

**Figure S5. Length distribution**, **nucleotide composition**, **and overlapping nucleotide distribution for piRNAs in the K63855 testis.**

**a.** All piRNA cluster-mapping piRNAs, corresponding to **Figure 2h**, **i**, **j** but for the oxidized K63464 testis.

**b.** Transposon-mapping piRNAs, corresponding to **Figure 3a**, **b**, **c** but for the oxidized K63464 testis.

**c.** Size distribution of piRNAs mapping to KoRV-A, Ko.ERV.1, Ko.ERVL.1, or Ko.ERVK.14, corresponding to **Figure 3d** but for oxidized piRNAs in the K63464 testis and unoxidized piRNAs in the K63464 and K63855 testis tissues, respectively.

**d.** Distributions of overlapping nucleotides between sense and antisense piRNAs from KoRV-A, Ko.ERV.1, Ko.ERVL.1, and Ko.ERVK.14, for oxidized piRNAs in the K63464 and K63855 testis and unoxidized piRNAs in the K63464 and K63855 testis tissues, respectively.

**Figure S6. piRNAs derived from genic piRNA clusters are preferentially spliced.**

**a.** Long RNA reads across KoRV-A in liver and brain, with the junction to splice-site ratios for the main exon-exon junction indicated. These figures correspond to **Figure 4a** but for long RNA reads in liver and brain samples of two Koalas.

**b.** Boxplots indicating splicing index of 2,460 exon-exon junctions (EEJ) in 204 genic piRNA clusters. Splicing index defined by long RNA reads is marked as green while splicing index defined by piRNA reads are marked as purple. We could only detect junction-or splice site-mapping piRNA reads and long RNA reads at 1,404 junctions.

**c.** Long RNA and piRNA reads across the fourth to eighth exons of the genic piRNA cluster pi-phaCin-PC-ARHGAP20 in testis of the koala K63464. Most long RNAs and piRNAs are spliced at the exon-exon junctions.

**d.** A scatterplot comparing the splicing index defined by long RNA reads and piRNA reads for all transposon exon-exon junctions in the mouse adult testis tissue. This panel corresponds to **Figure 4c**, which were for sorted mouse pachytene spermatocyte and round spermatid cells.

**e.** Long RNA and piRNA reads across the ERV1_MD_I transposon in opossum testis, with the ratio of junction reads over splice-site reads for the annotated exon-exon junction indicated. This figure corresponds to **Figure 4a** but for ERV1_MD_I in opossum.

**f.** Barplots showing splicing index of KoRV-A and Ko.ERV.1 transcripts in K63464 and K63855 testis, brain and liver tissues.

**Table S1. Detailed information on 379 piRNA clusters.**

**Table S2. Mapping statistics of DNA-seq**, **RNA-seq and small RNA-seq data analyzed in this paper.**

**Table S3. Summarized information for koala transposon consensus sequences and their copies in the koala reference genome.**

## METHODS

### Data availability

All data have been deposited to GEO with the accession number GSE128122.

### Experimental details

#### Total DNA/RNA isolation

Total DNA was isolated from koala (K63464 and K63855) tissue samples using the DNeasy® Blood and Tissue Kit (Qiagen). Total RNA was isolated from brain, liver, and testis from two koalas (K63464 and K63855) using the mirVana™ miRNA Isolation Kit (Life Technologies). Total RNA samples were treated with Turbo DNase (Invitrogen) and RNA cleanup was done following the manufacturer’s instructions using the RNeasy Mini Kit (Qiagen).

#### Library preparation

Small RNA sequencing libraries were prepared as previously described^42^. Briefly, total RNA was isolated from flash frozen koala tissue using the mirVana miRNA Isolation Kit (Ambion/Life Technologies). Small RNAs were sized selected and purified from a 15% denaturing polyacrylamide-urea gel using the ZR small-RNA™ PAGE Recovery Kit (Zymo Research). For oxidized RNA library preparations, purified RNA was oxidized with 25 mM NaIO4 in 30 mM borax, 30 mM boric acid, pH 8.6, for 30 min at room temperature followed by ethanol precipitation. 3’ pre-adenylated adapter was ligated to oxidized or un-oxidized small RNAs. The 3’ ligated product was purified from a 15% denaturing polyacrylamide-urea gel. 5’ RNA adapter ligation was performed and the ligated product was purified from a 10% denaturing polyacrylamide-urea gel and used to synthesize cDNA. The resulting cDNA was PCR amplified and run on a 2% Certified Low Range Ultra Agarose (Bio-Rad) gel with subsequent extraction using the QIAquick® Gel Extraction Kit (Qiagen). Purified small RNA libraries were single-end sequenced using the Illumina NextSeq 500 system.

Strand-specific RNA sequencing libraries for one K63464 testis replicate (rep2) and two K63855 testis replicates were prepared as previously described^43^. In brief, total RNA was isolated from frozen koala tissue samples via the mirVana miRNA Isolation kit and then rRNA depleted using the RiboZero™ Gold rRNA Removal Kit for human, mouse, and rat (Illumina). RNAs longer than 200 nt were selectively recovered using the RNA Clean & Concentrator-5 kit (Zymo Research). RNA samples were fragmented and reverse transcribed. dUTP was incorporated during second strand synthesis for strand specificity. End repair and A-tailing was performed followed by adapter ligation and uracil-DNA glycosylase (UDG) treatment. Finally, the library was PCR amplified. RNA libraries were paired-end sequenced using the Illumina NextSeq 500 system. Strand-specific RNA-seq libraries for one K63464 testis replicate (rep1), K63464 liver, K63464 brain, K63855 liver, and K63855 brain are constructed at BGI. Then these RNA-seq libraries are sequenced in HiSeq 101PE platform at BGI.

Short fragments of DNA libraries for K63464 testis, K63464 liver, K63464 brain, K63855 liver, and K63855 brain are constructed at BGI. Then these DNA-seq libraries were sequenced using the NovaSeq 151PE platform at BGI.

### Bioinformatics analysis

#### Transposon annotation

We annotated transposon consensus sequences and individual copies in the koala reference genome using several algorithms: repeatModeler, repeatMasker, LTRharvest, LTRdigest, TransposonPSI, and ucluster^44–48^, described in detail as follows.

First, we used three separate algorithms to identify transposons de novo. We ran RepeatModeler on the koala reference genome with default parameters to build a library of transposons. We also used LTRharvest with parameters “-index phaCin.fsa -seed 100 -minlenltr 100 -maxlenltr 1000 - mindistltr 1000 -maxdistltr 15000 -xdrop 5 -mat 2 -mis -2 -ins -3 -del -3 -similar 90.0 -overlaps best-mintsd 5 -maxtsd 20 -motif tgca -motifmis 0 -vic 60 -longoutput” to predict LTR retrotransposons. LTRdigest was then used to filter out false positives from LTR transposons predicted by LTRharvest using the hidden Markov models (HMMs) of transposon proteins from Pfam and GyDB. LTR elements without HMM hits were discarded. Finally, transposonPSI was used to construct all potential transposon sequences based on high homology to existing transposon annotation.

Second, we merged the three transposon libraries by RepeatModeler, LTRharvest/LTRdigest, and TransposonPSI via usearch (version 11.0.667). Transposon clusters with fewer than 3 transposon sequences were discarded. The remaining transposon clusters were deemed de novo discovered transposon families with their centroid sequences regarded as the consensus sequences. We further combined the de novo transposon library with Repbase’s transposon library of koala via usearch and provided the centroid sequences to RepeatClassfier to determine their transposon families, and only those transposons classified as LINE/SINE/LTR/DNA were retained in our final consensus sequence library.

Finally, we provided our final consensus sequences to RepeatMasker with parameters “-s -pa 48 - e ncbi -div 40 -nolow -norna -no_is” to identify individual copies in the genome. To further remove false positives, we discarded those transposon families without at least two copies or 1000 base pairs in the koala reference genome.

Except for KoRV-A and PhER which had been named already, we assigned transposons names that began with Ko, which stood for koala, followed by the family they belonged to. A number was also added for different subfamilies in the same family. For example, ERV family transposons were named Ko.ERV.1, Ko.ERV.2, Ko.ERV.3, etc.

#### Calculation of average sequence divergence for transposon subfamilies

For each transposon subfamily, all copies annotated in reference genome were extracted, and copies longer than half of the length of the consensus sequence were aligned to the consensus sequence, and the alignments were used to calculate the average divergence for each transposon subfamily, defined as the number of substituted nucleotides in the alignments divided by the total number of nucleotides of these copies in the reference genome.

#### Quantification of the expression levels of transposons and piRNA pathway genes

We first removed all RNA-seq reads that mapped to ribosomal RNAs (rRNAs) in the reference koala genome. We annotated the rRNAs using RNAmmer^49,50^ (version 1.2) with default parameters and mapped RNA-seq reads to rRNAs using Bowtie2^51,52^ (version 2.2.5) with default parameters. The remaining RNA-seq reads were mapped to the reference koala genome using STAR^51^ (version 020201). The mapping results in the SAM format were transformed into sorted and duplication-removed BAM format using SAMtools (version 1.8). The final mapped reads were assigned to protein-coding genes, non-coding RNAs, and piRNA genes using HTSeq^53^ (version 0.9.1), and the expression levels of these genes, in reads per million unique mapped reads in per thousand nucleotides (RPKM), were calculated using custom bash scripts.

After rRNA removal, RNA-seq reads were also mapped to transposon consensus sequences (defined as described in the previous sub-section) directly using Hisat2^53,54^ (version 2.1.0) with default parameters. Then transposon expression levels were calculated using Bedtools^55^ (version 2.27.1). The similarity between some transposon elements may prevent the reads from these high-similarity regions to be counted accurately when these reads are mapped directly to the transposon consensus sequences. Thus, we applied the expectation-maximization (EM) algorithm, which assigns these multiple mappers according to mapping potential in different transposons^56^.First, multiple mappers were assigned to different transposons according to the number of unique mappers in each transposons. Then the number of assigned reads for each transposon was used as the weight for the second round of multiple mapper assignments. The iterations continued until the process converged, defined as the Manhattan distance between two iterations being less than 0.05.

#### Identification of new transposon insertions and transposon absences using genomic DNA-seq data

DNA-seq raw reads were mapped to the reference koala genome using the BWA MEM algorithm^57^ (version 0.7.12-r1044) allowing soft clipping. Mapping results in the SAM format were transformed into sorted BAM format via SAMtools^49^ (version 1.8). We then used the TEMP algorithm^58^ to detect new transposon insertions. TEMP defines discordant read pairs as those with one end mapping to the reference genome and the other mapping to an inserted transposon element and uses these discordant reads for identifying new transposon insertions. Such discordant reads may also come from low-frequency chimeric reads that result from sequencing library construction^25^, but in such case, the transposon mapping ends would be randomly distributed and randomly oriented over the transposon element. Thus, we only retained reads mapping to the ends of the element, with the correct orientation for a bona fide insertion. We made two more modifications to TEMP for detecting transposons that are in the reference genome but not in the new sample. First, we only used reads flanking ±5 bp of breakpoints as supporting reads instead of the default 0–5 bp in TEMP. Second, the default TEMP uses all reads overlapped with the region from upstream 20 bp of the 5’-end to downstream 20 bp of the 3’-end of a transposon insertion to detect breakpoints. Instead, we only used reads that overlapped with the two ends of a transposon insertion, ±20 bp centered on the 5’-end or ±20 bp centered on the 3’-end, to detect breakpoints, which is more stringent and guards against low-frequency chimeric reads that result from sequencing library construction^25^.

#### Analysis of small RNA-seq data

We first removed the adapter sequence (TGGAATTCTCGGGTGCCAAGGAACTCCAGTCACCGATGTATCTCGT) from all small RNA-seq reads using cutadapt^59^ (version 1.15). We then removed all reads that mapped to rRNAs, miRNAs, snoRNAs, snRNAs, and tRNAs. As mentioned above, we defined rRNAs in the koala genome using RNAmmer^49,50^ (version 1.2) with default parameters. We annotated miRNAs in the koala genome using miRDeep2^60^ (version 2.0.0.8) with default parameters and the unoxidized small RNA-seq datasets in the testis of K63464 and K63855 testes as the input. The remaining small RNA-seq reads were mapped to reference koala genome and transposon consensus sequences independently using Bowtie (version 1.1.0). We calculated piRNA abundance at piRNA clusters and individual transposon insertions in the genome using unoxidized and oxidized small RNA-seq samples separately. We normalized the abundance by sequencing depth, i.e., the total number of genome mapping reads after removing rRNAs, miRNAs, snoRNAs, snRNAs, and tRNAs. Nucleotide content and ping-pong amplification were analyzed for all reads that mapped to the genome, reads that mapped to transposons, and reads that mapped to piRNA clusters, respectively. For ping-pong analysis on cluster-mapping reads, 5’ to 5’ overlaps between all pairs of piRNAs that mapped to the opposite genomic strands were calculated, and then the Z-score for the 10-nt overlap was calculated using the 1-9 nt and 11-30 nt overlaps as the background^42^. For ping-pong analysis on transposon-mapping reads, we only used the reads that mapped to transposon consensus sequences with zero or one mismatch and the 10-nt overlap was computed using the coordinates of reads on the consensus sequences. Reads that mapped to the Ko.RTEBovB.98 transposon were excluded due to one highly abundant sequence.

#### piRNA cluster annotation

We annotated piRNA clusters using RNA-seq and small RNA-seq data in the K63464 testis. We considered 24–32 nt small RNA reads that could map to the koala genome, after rRNA, miRNA, tRNA, snRNA, and snoRNA were removed, as piRNAs. piRNAs were then assigned to 20 kb sliding windows (with a 1 kb step), and windows with more than 100 piRNAs per million uniquely mapped piRNAs were further considered as potential piRNA clusters. To remove false positives due to un-annotated miRNA, rRNA, tRNA, snRNA, and snoRNA, which mostly produce reads with the same sequences, we also filtered out those 20-kb genomic windows with fewer than 200 distinct reads (called species). We then calculated the first-nucleotide content for each 20-kb window, and those windows with 1U/10A percentage less than 50% were also discarded (15 windows were discarded in total). The remaining contiguous 20-kb windows were deemed putative piRNA clusters. We used the RNA-seq reads after rRNA removal for annotating exon-exon junctions in piRNA clusters. Finally, we performed manual curation for putative piRNA clusters using piRNA profile, long RNA profile, and detected exon-exon junctions. The direction of a piRNA cluster was indicated by the direction of the main long-RNA transcript.

The final piRNA clusters were classified according to their genomic location, directionality, and strandedness. First, piRNA clusters with more than 50% base pairs overlapped with protein-coding genes were defined as genic piRNA clusters and others intergenic piRNA clusters. Second, piRNA cluster pairs that shared their promoters (distance between their TSSs were less than 500 bp) but transcribe in divergent directions were annotated as bidirectional piRNA clusters while others were unidirectional piRNA clusters. Third, piRNA clusters which produced similar levels of piRNAs from both genomic strands (< 4 fold difference) were defined as dual-strand piRNA clusters while others were uni-strand clusters.

#### Exon-exon junction discovery and splicing index calculation in transposons

RNA-seq data (after rRNA removal) were used to identify high-confident exon-exon junctions in transposons. First, RNA-seq reads that could map fully to the reference koala genome (using Bowtie2 in the end-to-end mode and soft clipping disabled) were considered primary transcripts and discarded. The remaining RNA-seq reads were then mapped to the transposon consensus sequences using Hisat2^54^ with default parameters for exon-exon junction detection.

To calculate the splicing index for each detected junction, rRNA-removed long RNA reads and rRNA/miRNA/tRNA/snRNA/snoRNA removed small RNA reads were used as input, and the calculation was performed in the following steps. First, reads were mapped to transposon consensus sequences using Hisat2 given the detected junctions. The output of Hisat2 contained two types of reads, those that mapped to splice sites (unspliced reads) and the remaining that mapped to exon-exon junctions. We further mapped the junction-mapping reads back to the reference genome (Bowtie2 in the end-to-end mode with soft clipping disabled) and discarded the reads that could map. The remaining long RNA reads that spanned at least 7 bps of the exon-exon junctions and piRNA reads that spanned at least 3 bps of the exon-exon junctions were considered spliced reads. After this, splicing index for each exon-exon junction detected by long RNA and piRNA respectively was defined as the ratio of spliced reads and the total reads (splice reads plus unspliced reads).

## Acknowledgments

We would like to thank the members of Theurkauf and Weng laboratories for their critical comments and discussion on the manuscript. This work was supported in part by NIH grants HD078253 and HD049116 (to W.T. and Z.W.) and Chinese National Natural Science Foundation grants (31571362 and 31871296, and 91640201 to Z.W.).

## Author contributions

T.Y., B.S.K., Z.W., and W.E.T. conceived the project. B.S.K. performed the experiments. T.Y. performed the computational analyses. K.C. and S.J. provided the tissues of two koalas. S.P. performed the dissection and freezing of the koala tissues. T.Y., Z.W., and W.E.T. wrote the manuscript.

### Correspondence author

Correspondence should be addressed to William E. Theurkauf or Zhiping Weng.

## References

1. Iskow, R. C. et al. Natural mutagenesis of human genomes by endogenous retrotransposons. Cell 141, 1253–1261 (2010).

2. Belancio, V. P., Hedges, D. J. & Deininger, P. Mammalian non-LTR retrotransposons: for better or worse, in sickness and in health. Genome Res. 18, 343–358 (2008).

3. Aravin, A. A., Hannon, G. J. & Brennecke, J. The Piwi-piRNA pathway provides an adaptive defense in the transposon arms race. Science 318, 761–764 (2007).

4. Denner, J. & Young, P. R. Koala retroviruses: characterization and impact on the life of koalas. Retrovirology 10, 108 (2013).

5. Tarlinton, R. E., Meers, J. & Young, P. R. Retroviral invasion of the koala genome. Nature 442, 79–81 (2006).

6. Weick, E.-M. -M. Weick, E. & Miska, E. A. piRNAs: from biogenesis to function. Development 141, 3458–3471 (2014).

7. Ernst, C., Odom, D. T. & Kutter, C. The emergence of piRNAs against transposon invasion to preserve mammalian genome integrity. Nat. Commun. 8, 1411 (2017).

8. Saito, K. et al. Specific association of Piwi with rasiRNAs derived from retrotransposon and heterochromatic regions in the Drosophila genome. Genes Dev. 20, 2214–2222 (2006).

9. Brennecke, J. et al. Discrete small RNA-generating loci as master regulators of transposon activity in Drosophila. Cell 128, 1089–1103 (2007).

10. Bergman, C. M., Quesneville, H., Anxolabéhère, D. & Ashburner, M. Recurrent insertion and duplication generate networks of transposable element sequences in the Drosophila melanogaster genome. Genome Biol. 7, R112 (2006).

11. Siomi, M. C., Sato, K., Pezic, D. & Aravin, A. A. PIWI-interacting small RNAs: the vanguard of genome defence. Nat. Rev. Mol. Cell Biol. 12, 246–258 (2011).

12. Mével-Ninio, M., Pelisson, A., Kinder, J., Campos, A. R. & Bucheton, A. The flamenco locus controls the gypsy and ZAM retroviruses and is required for Drosophila oogenesis. Genetics 175, 1615–1624 (2007).

13. Sarot, E., Payen-Groschêne, G., Bucheton, A. & Pélisson, A. Evidence for a piwi-dependent RNA silencing of the gypsy endogenous retrovirus by the Drosophila melanogaster flamenco gene. Genetics 166, 1313–1321 (2004).

14. Li, X. Z. et al. An ancient transcription factor initiates the burst of piRNA production during early meiosis in mouse testes. Mol. Cell 50, 67–81 (2013).

15. Wasik, K. A. et al. RNF17 blocks promiscuous activity of PIWI proteins in mouse testes. Genes Dev. 29, 1403–1415 (2015).

16. Rio, D. Regulation of Drosophila P element transposition. Trends in Genetics 7, 282–287 (1991).

17. Khurana, J. S. et al. Adaptation to P Element Transposon Invasion in Drosophila melanogaster. Cell 147, 1551–1563 (2011).

18. Brennecke, J. et al. An epigenetic role for maternally inherited piRNAs in transposon silencing. Science 322, 1387–1392 (2008).

19. Chappell, K. J. et al. Phylogenetic Diversity of Koala Retrovirus within a Wild Koala Population. J. Virol. 91, (2017).

20. Ishida, Y., Zhao, K., Greenwood, A. D. & Roca, A. L. Proliferation of endogenous retroviruses in the early stages of a host germ line invasion. Mol. Biol. Evol. 32, 109–120 (2015).

21. Johnson, R. N. et al. Adaptation and conservation insights from the koala genome. Nat. Genet. 50, 1102–1111 (2018).

22. Ávila-Arcos, M. C. et al. One hundred twenty years of koala retrovirus evolution determined from museum skins. Mol. Biol. Evol. 30, 299–304 (2013).

23. Hobbs, M. et al. A transcriptome resource for the koala (Phascolarctos cinereus): insights into koala retrovirus transcription and sequence diversity. BMC Genomics 15, 786 (2014).

24. Cui, P. et al. Comprehensive profiling of retroviral integration sites using target enrichment methods from historical koala samples without an assembled reference genome. PeerJ 4, e1847 (2016).

25. Treiber, C. D. & Waddell, S. Resolving the prevalence of somatic transposition in. Elife 6, (2017).

26. Gunawardane, L. S. et al. A slicer-mediated mechanism for repeat-associated siRNA 5’ end formation in Drosophila. Science 315, 1587–1590 (2007).

27. Horwich, M. D. et al. The Drosophila RNA methyltransferase, DmHen1, modifies germline piRNAs and single-stranded siRNAs in RISC. Curr. Biol. 17, 1265–1272 (2007).

28. Saito, K. et al. Pimet, the Drosophila homolog of HEN1, mediates 2’-O-methylation of Piwi-interacting RNAs at their 3’ ends. Genes Dev. 21, 1603–1608 (2007).

29. Aravin, A. A., Sachidanandam, R., Girard, A., Fejes-Toth, K. & Hannon, G. J. Developmentally Regulated piRNA Clusters Implicate MILI in Transposon Control. Science 316, 744–747 (2007).

30. Aravin, A. A. et al. A piRNA pathway primed by individual transposons is linked to de novo DNA methylation in mice. Mol. Cell 31, 785–799 (2008).

31. Zanni, V. et al. Distribution, evolution, and diversity of retrotransposons at the flamenco locus reflect the regulatory properties of piRNA clusters. Proc. Natl. Acad. Sci. U. S. A. 110, 19842– 19847 (2013).

32. Paul, W. E. Bridging Innate and Adaptive Immunity. Cell 147, 1212–1215 (2011).

33. Dumesic, P. A. et al. Stalled spliceosomes are a signal for RNAi-mediated genome defense. Cell 152, 957–968 (2013).

34. Zhang, G. et al. Co-dependent Assembly of Drosophila piRNA Precursor Complexes and piRNA Cluster Heterochromatin. Cell Rep. 24, 3413–3422.e4 (2018).

35. Sakuma, T., Tonne, J. M. & Ikeda, Y. Murine leukemia virus uses TREX components for efficient nuclear export of unspliced viral transcripts. Viruses 6, 1135–1148 (2014).

36. Bradshaw, V. A. & McEntee, K. DNA damage activates transcription and transposition of yeast Ty retrotransposons. Mol. Gen. Genet. 218, 465–474 (1989).

37. Farkash, E. A. Gamma radiation increases endonuclease-dependent L1 retrotransposition in a cultured cell assay. Nucleic Acids Research 34, 1196–1204 (2006).

38. McClintock, B. The significance of responses of the genome to challenge. Science 226, 792– 801 (1984).

39. Fiebig, U., Hartmann, M. G., Bannert, N., Kurth, R. & Denner, J. Transspecies transmission of the endogenous koala retrovirus. J. Virol. 80, 5651–5654 (2006).

40. Brubaker, S. W., Bonham, K. S., Zanoni, I. & Kagan, J. C. Innate immune pattern recognition: a cell biological perspective. Annu. Rev. Immunol. 33, 257–290 (2015).

41. Bonilla, F. A. & Oettgen, H. C. Adaptive immunity. J. Allergy Clin. Immunol. 125, S33–40 (2010).

42. Li, C. et al. Collapse of germline piRNAs in the absence of Argonaute3 reveals somatic piRNAs in flies. Cell 137, 509–521 (2009).

43. Zhang, Z., Theurkauf, W. E., Weng, Z. & Zamore, P. D. Strand-specific libraries for high throughput RNA sequencing (RNA-Seq) prepared without poly(A) selection. Silence 3, 9 (2012).

44. Website. Available at: .“> Smit, AFA, Hubley, R & Green, P. RepeatMasker Open-4.0. 2013-2015 http://www.repeatmasker.org. (Accessed: 31st December 2018)

45. Ellinghaus, D., Kurtz, S. & Willhoeft, U. LTRharvest, an efficient and flexible software for de novo detection of LTR retrotransposons. BMC Bioinformatics 9, 18 (2008).

46. Steinbiss, S., Willhoeft, U., Gremme, G. & Kurtz, S. Fine-grained annotation and classification of de novo predicted LTR retrotransposons. Nucleic Acids Res. 37, 7002–7013 (2009).

47. Website. Available at: TransposonPSI. http://transposonpsi.sourceforge.net. (Accessed: 31st December 2018)

48. Edgar, R. C. Search and clustering orders of magnitude faster than BLAST. Bioinformatics 26, 2460–2461 (2010).

49. Li, H. et al. The Sequence Alignment/Map format and SAMtools. Bioinformatics 25, 2078– 2079 (2009).

50. Lagesen, K. et al. RNAmmer: consistent and rapid annotation of ribosomal RNA genes. Nucleic Acids Res. 35, 3100–3108 (2007).

51. Dobin, A. et al. STAR: ultrafast universal RNA-seq aligner. Bioinformatics 29, 15–21 (2013).

52. Langmead, B. & Salzberg, S. L. Fast gapped-read alignment with Bowtie 2. Nat. Methods 9, 357–359 (2012).

53. Anders, S., Pyl, P. T. & Huber, W. HTSeq--a Python framework to work with high-throughput sequencing data. Bioinformatics 31, 166–169 (2015).

54. Kim, D., Langmead, B. & Salzberg, S. L. HISAT: a fast spliced aligner with low memory requirements. Nat. Methods 12, 357–360 (2015).

55. Malloy, A. A benchmarking suite for comparing middleware performance. doi:10.22215/etd/2011-08952

56. Fu, Y. et al. The genome of the Hi5 germ cell line from Trichoplusia ni, an agricultural pest and novel model for small RNA biology. Elife 7, (2018).

57. Li, H. & Durbin, R. Fast and accurate short read alignment with Burrows-Wheeler transform. Bioinformatics 25, 1754–1760 (2009).

58. Zhuang, J., Wang, J., Theurkauf, W. & Weng, Z. TEMP: a computational method for analyzing transposable element polymorphism in populations. Nucleic Acids Res. 42, 6826–6838 (2014).

59. Martin, M. Cutadapt removes adapter sequences from high-throughput sequencing reads. EMBnet.journal 17, 10 (2011).

60. Friedländer, M. R., Mackowiak, S. D., Li, N., Chen, W. & Rajewsky, N. miRDeep2 accurately identifies known and hundreds of novel microRNA genes in seven animal clades. Nucleic Acids Res. 40, 37–52 (2012).

