## Supplemental figures for "The piRNA Response to Retroviral Invasion of the Koala Genome"

Yu et al., Figure S1

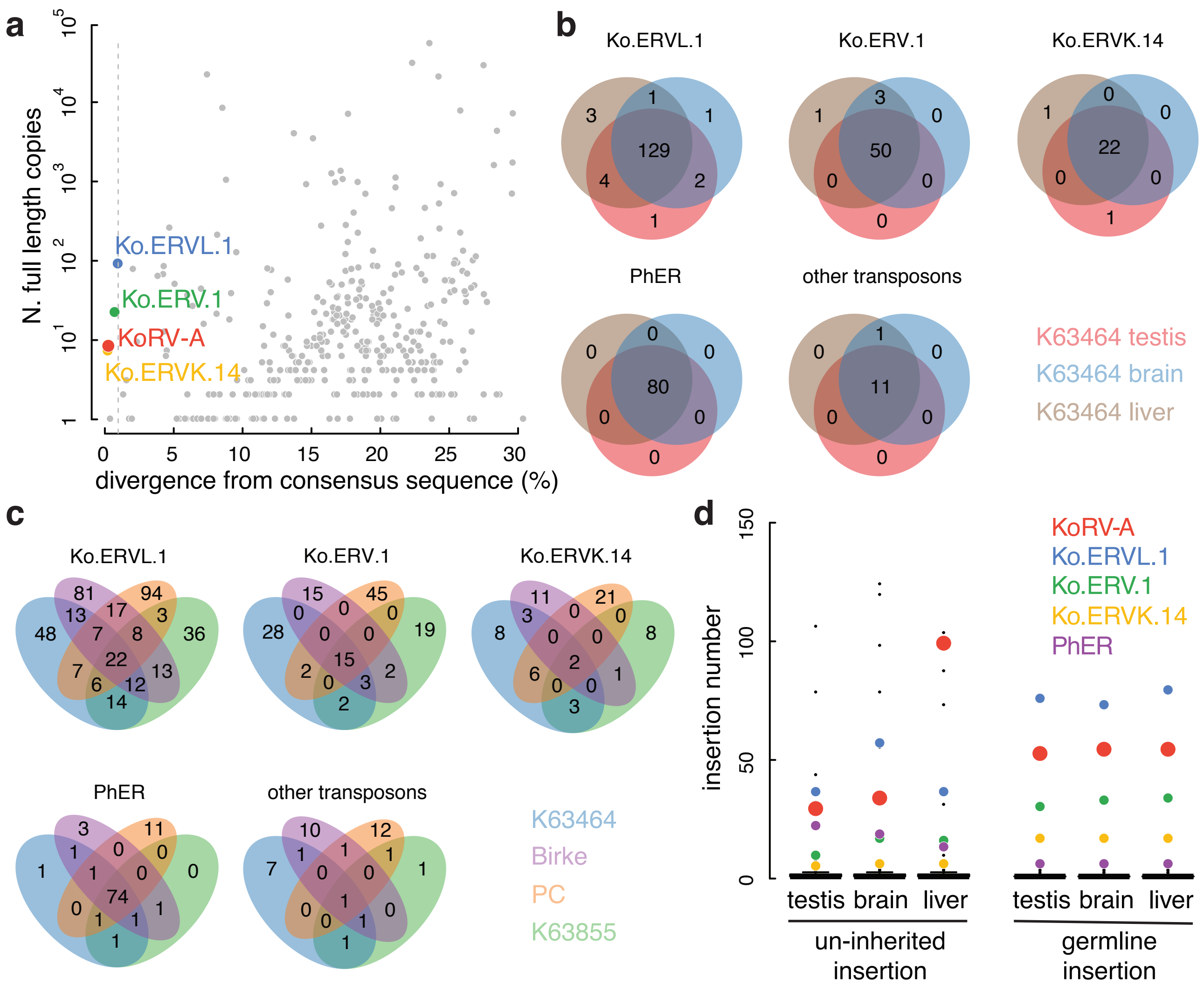

Yu et al., Figure S2

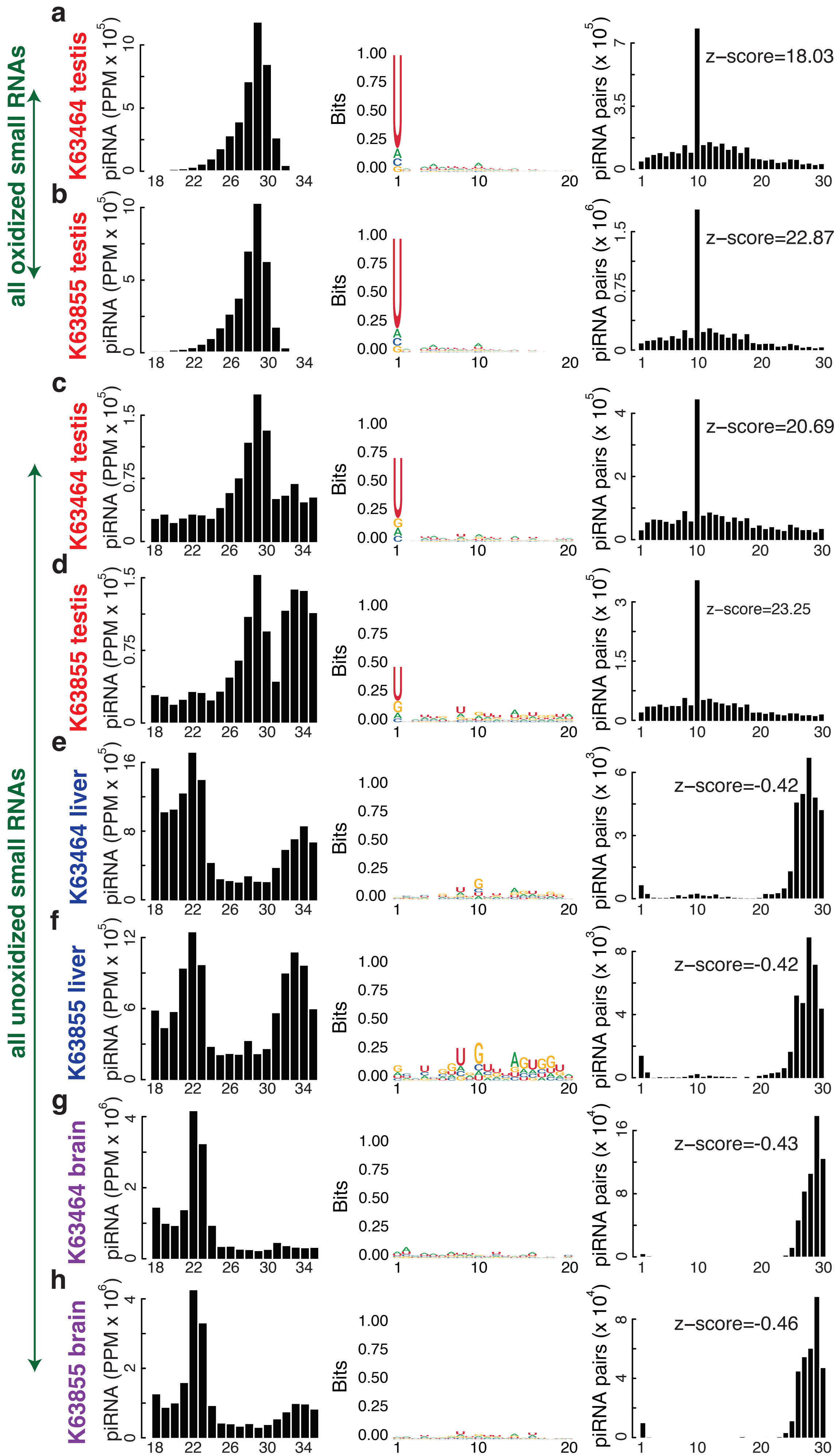

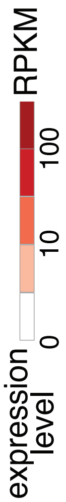


| | K63464 testis | K63464 brain | K63464 liver | K63855 testis | K63855 brain | K63855 liver | BI testis | BI boneMarrow | BI kidney | BI liver | BI lymphNode | BI salivaryGland | BI spleen | PC uterus | PC adrenal | PC brain | PC heart | PC kidney | PC liver | PC lung | PC spleen |
|---|---|---|---|---|---|---|---|---|---|---|---|---|---|---|---|---|---|---|---|---|---|
| *PIWIL1* | 2.23 | 0.36 | 0.00 | 1.61 | 0.11 | 0.00 | 1.72 | 0.00 | 0.00 | 0.00 | 0.00 | 0.00 | 0.00 | 0.08 | 0.00 | 0.00 | 0.00 | 0.00 | 0.00 | 0.02 | 0.00 |
| *PIWIL2* | 17.14 | 1.07 | 0.41 | 18.30 | 0.99 | 0.55 | 21.79 | 0.00 | 0.00 | 0.57 | 2.23 | 0.76 | 2.39 | 1.61 | 0.00 | 0.00 | 0.00 | 0.00 | 0.52 | 1.12 | 1.80 |
| *PIWIL4* | 4.89 | 0.00 | 0.00 | 3.34 | 0.00 | 0.00 | 5.23 | 0.00 | 0.00 | 0.00 | 0.01 | 0.00 | 0.00 | 0.01 | 0.00 | 0.00 | 0.00 | 0.00 | 0.00 | 0.02 | 0.05 |
| *MYBL1* | 15.52 | 0.18 | 0.01 | 14.36 | 0.12 | 0.04 | 9.09 | 0.00 | 0.00 | 0.02 | 6.68 | 0.06 | 1.76 | 0.10 | 0.00 | 0.00 | 0.00 | 0.00 | 0.02 | 0.19 | 1.36 |
| *TDRD1* | 19.48 | 0.02 | 0.00 | 14.23 | 0.00 | 0.00 | 7.79 | 0.00 | 0.00 | 0.00 | 0.01 | 0.01 | 0.00 | 0.00 | 0.00 | 0.00 | 0.00 | 0.00 | 0.00 | 0.18 | 0.00 |
| *TDRD5* | 37.46 | 0.19 | 2.93 | 29.29 | 0.15 | 2.75 | 25.80 | 0.00 | 0.00 | 3.05 | 0.65 | 0.21 | 0.25 | 0.08 | 0.00 | 0.00 | 0.00 | 0.00 | 2.94 | 0.17 | 0.27 |
| *TDRD6* | 9.73 | 0.25 | 0.01 | 8.16 | 0.13 | 0.02 | 4.66 | 0.00 | 0.00 | 0.01 | 0.05 | 0.01 | 0.08 | 0.00 | 0.00 | 0.00 | 0.00 | 0.00 | 0.02 | 0.09 | 0.01 |
| *TDRD12* | 11.60 | 1.94 | 0.51 | 9.62 | 2.08 | 0.45 | 3.49 | 0.00 | 0.00 | 0.21 | 0.88 | 1.01 | 1.33 | 2.28 | 0.00 | 0.00 | 0.00 | 0.00 | 0.94 | 2.28 | 2.37 |
| *PLD6* | 16.33 | 0.71 | 0.24 | 12.11 | 0.90 | 0.73 | 14.80 | 0.00 | 0.00 | 0.02 | 0.25 | 0.27 | 0.33 | 0.54 | 0.00 | 0.00 | 0.00 | 0.00 | 0.23 | 0.82 | 0.70 |
| *DDX39A* | 9.62 | 9.81 | 11.97 | 10.86 | 14.13 | 36.49 | 17.12 | 0.00 | 0.00 | 27.16 | 48.23 | 9.06 | 54.72 | 24.21 | 0.00 | 0.00 | 0.00 | 0.00 | 11.14 | 25.71 | 47.97 |
| *DDX39B* | 22.65 | 54.46 | 58.48 | 24.11 | 58.58 | 63.06 | 73.82 | 0.00 | 0.00 | 38.02 | 80.61 | 50.46 | 72.49 | 72.82 | 0.00 | 0.00 | 0.00 | 0.00 | 54.31 | 82.57 | 122.42 |
| *MAEL* | 8.85 | 0.32 | 0.09 | 6.84 | 0.22 | 0.00 | 11.53 | 0.00 | 0.00 | 0.00 | 0.03 | 0.00 | 0.04 | 0.26 | 0.00 | 0.00 | 0.00 | 0.00 | 0.10 | 0.50 | 0.02 |
| *TDRD9* | 14.15 | 0.35 | 2.46 | 12.25 | 1.30 | 2.12 | 9.86 | 0.00 | 0.00 | 1.12 | 0.93 | 0.33 | 1.78 | 0.02 | 0.00 | 0.00 | 0.00 | 0.00 | 0.48 | 0.05 | 0.79 |
| *GTSF1* | 87.82 | 1.55 | 32.24 | 58.51 | 1.81 | 49.95 | 96.77 | 0.00 | 0.00 | 45.48 | 45.95 | 9.92 | 28.34 | 3.65 | 0.00 | 0.00 | 0.00 | 0.00 | 49.88 | 37.69 | 34.15 |
| *MOV10L1* | 23.13 | 4.28 | 5.28 | 15.18 | 3.31 | 0.43 | 18.94 | 0.00 | 0.00 | 1.33 | 2.28 | 0.10 | 0.54 | 0.26 | 0.00 | 0.00 | 0.00 | 0.00 | 1.75 | 0.44 | 0.51 |
| *GPAT2* | 3.55 | 0.50 | 0.01 | 1.83 | 0.21 | 0.00 | 6.69 | 0.00 | 0.00 | 0.00 | 0.12 | 0.80 | 0.55 | 0.01 | 0.00 | 0.00 | 0.00 | 0.00 | 0.02 | 0.37 | 0.22 |
| *DDX4* | 51.54 | 0.07 | 0.17 | 40.88 | 0.01 | 0.05 | 31.78 | 0.00 | 0.00 | 0.02 | 9.45 | 0.15 | 0.69 | 0.39 | 0.00 | 0.00 | 0.00 | 0.00 | 0.06 | 0.66 | 0.61 |
| *RNF17* | 42.16 | 0.10 | 0.04 | 35.91 | 0.14 | 0.22 | 21.94 | 0.00 | 0.00 | 0.36 | 0.53 | 0.05 | 0.51 | 0.37 | 0.00 | 0.00 | 0.00 | 0.00 | 0.28 | 0.30 | 0.13 |
| *TCFL5* | 21.78 | 1.43 | 0.23 | 19.56 | 1.33 | 0.13 | 27.28 | 0.00 | 0.00 | 0.17 | 1.14 | 0.90 | 0.83 | 0.85 | 0.00 | 0.00 | 0.00 | 0.00 | 0.19 | 0.63 | 0.66 |

Yu et al., Figure S4

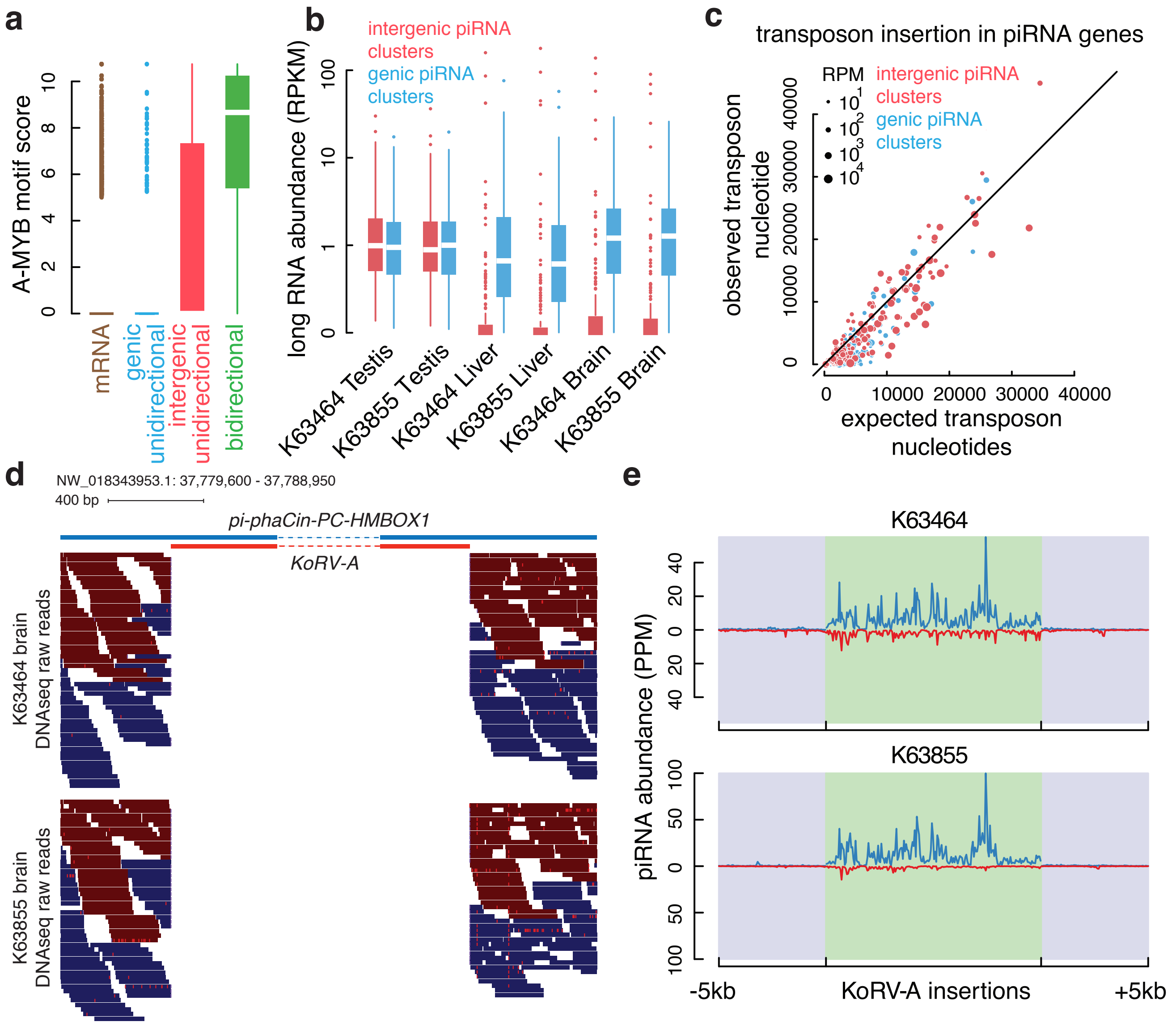

Yu et al., Figure S5

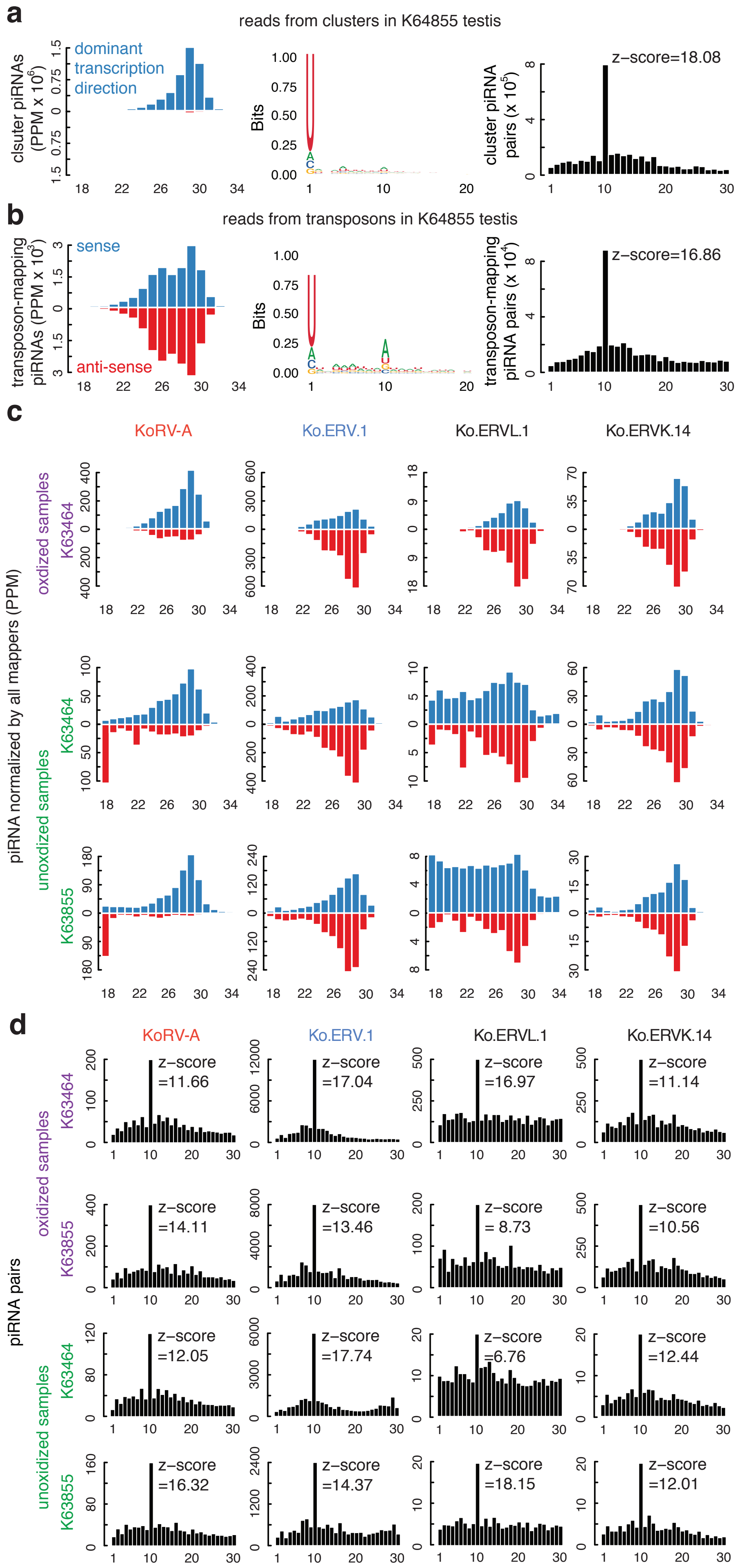

Yu et al., Figure S6

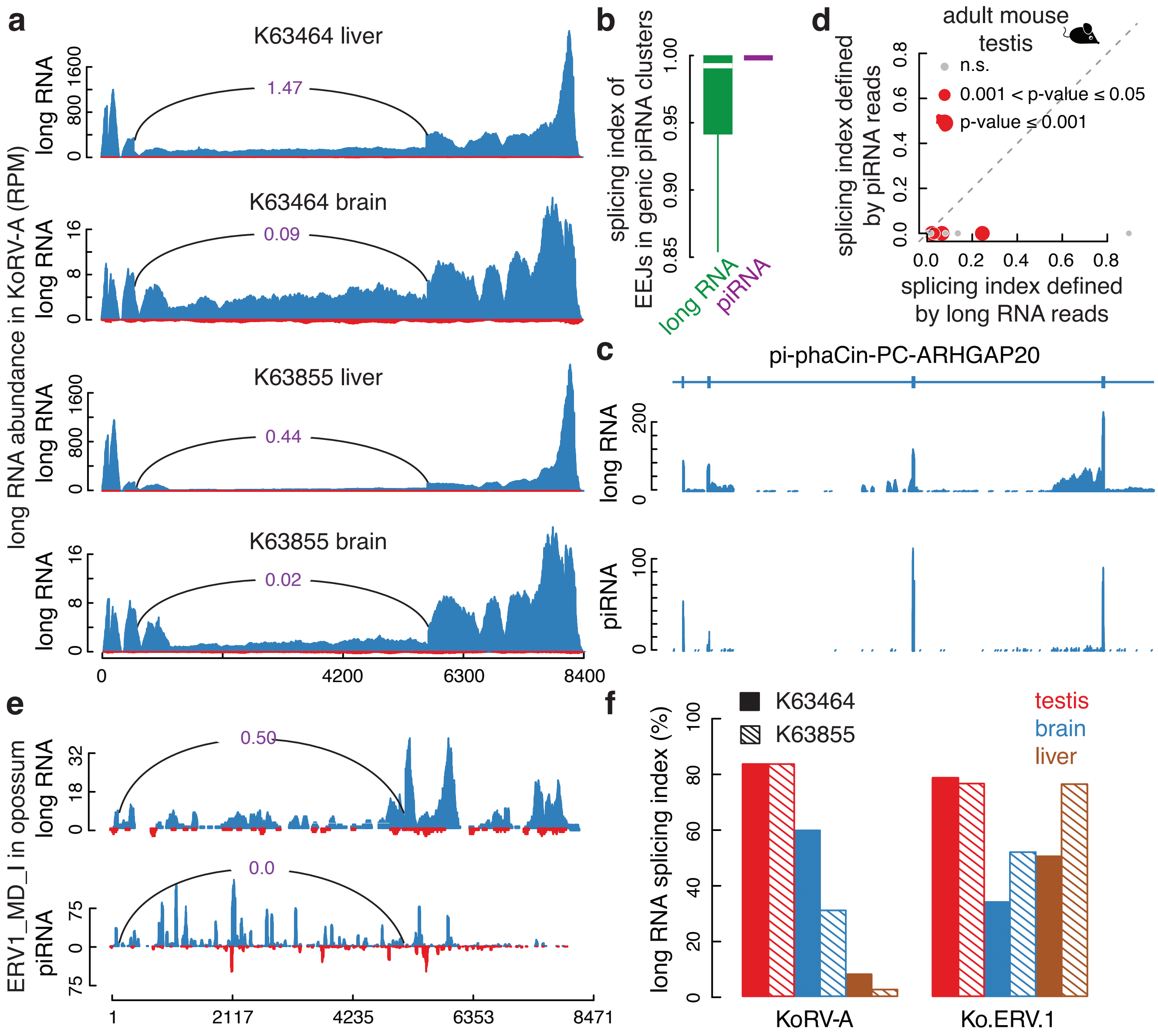
